# TMS cortical mapping of multiple muscles: absolute and relative test-retest reliability

**DOI:** 10.1101/2020.09.15.298794

**Authors:** Maria Nazarova, Pavel Novikov, Ekaterina Ivanina, Ksenia Kozlova, Larisa Dobrynina, Vadim V. Nikulin

## Abstract

The spatial accuracy of TMS may be as small as a few millimeters. Despite such great potential, navigated TMS (nTMS) mapping is still underused for the assessment of motor plasticity, particularly in clinical settings. Here we investigate the within-limb somatotopy gradient as well as absolute and relative reliability of three hand muscle cortical representations (MCRs) using a comprehensive grid-based sulcus-informed nTMS motor mapping. We enrolled 22 young healthy male volunteers. Two nTMS mapping sessions were separated by 5-10 days. Motor evoked potentials were obtained from abductor pollicis brevis (APB), abductor digiti minimi, and extensor digitorum communis. In addition to individual MRI-based analysis, we studied MNI normalized MCRs. For the reliability assessment, we calculated intra-class correlation and the smallest detectable change. Our results revealed a somatotopy gradient reflected by APB MCR having the most lateral location. Reliability analysis showed that the commonly used metrics of MCRs, such as areas, volumes, centers of gravity (COGs), and hotspots had a high relative and low absolute reliability for all three muscles. For within-limb TMS somatotopy, the most common metrics such as the shifts between MCR COGs and hotspots had poor relative reliability. However, overlaps between different muscle MCRs were highly reliable. We thus provide novel evidence that inter-muscle MCR interaction can be reliably traced using MCR overlaps while shifts between the COGs and hotspots of different MCRs are not suitable for this purpose. Our results have implications for the interpretation of nTMS motor mapping results in healthy subjects and patients with neurological conditions.

## 1. Introduction

Transcranial magnetic stimulation (TMS) is one of the main techniques for the non-invasive investigation of the motor-cortex in humans. Measuring the effects of the multiple spot stimulation is a powerful approach for brain mapping, especially when it is combined with MRI navigation (called navigated TMS or nTMS) – a procedure which has been FDA approved for the presurgical brain mapping since 2009 (Krieg, 2017). Recently, nTMS motor mapping has been demonstrated to be even more accurate than functional MRI (fMRI) motor mapping, when compared with the direct cortical stimulation results (Weiss Lucas et al., 2020). Most commonly, during TMS motor mapping one investigates motor evoked potentials (MEPs) using surface (Rossini et al., 2015) or needle EMG (Massé-Alarie, Bergin, Schneider, Schabrun, & Hodges, 2017). When MEPs from the stimulation of many cortical points are acquired, the resulting output is referred to as cortical muscle representation (MCR) (Bashir, Perez, Horvath, & Pascual-Leone, 2013; de Carvalho, Miranda, Luis, & Ducla-Soares, 1999), also known as TMS cortical motor map (Kraus & Gharabaghi, 2015; Novikov, Nazarova, & Nikulin, 2018). There are numerous studies showing that the MCR parameters such as excitability, size and topography reflect functionally relevant features of the motor cortex organization in healthy people (Beaulieu, Flamand, Massé-Alarie, & Schneider, 2017; Gentner & Classen, 2006; Nazarova, Novikov, Nikulin, & Ivanova, 2020; Tyč & Boyadjian, 2011) and in patients with motor pathology such as stroke (Lüdemann-Podubecká & Nowak, 2016; Yarossi et al., 2019), amyotrophic lateral sclerosis (Chervyakov et al., 2015; de Carvalho et al., 1999), dystonia (Schabrun, Stinear, Byblow, & Ridding, 2009) etc. Considering the nTMS mapping of multiple muscles, the existence of the somatotopy gradient for MCRs, along with their extensive overlap, has been discussed from the earliest TMS studies (Gentner & Classen, 2006; Metman, Bellevich, Jones, Barber, & Streletz, 1993). The spatial accuracy of TMS, depending on stimulation parameters, based on animal data may be as small as 2 mm (Romero, Davare, Armendariz, & Janssen, 2019), thus allowing the tracing of within-limb MCR interactions, especially when taking into account individual sulcus anatomy (Raffin, Pellegrino, Di Lazzaro, Thielscher, & Siebner, 2015). These multiple-muscle MCR interactions are believed to reflect the basic neuroanatomical prerequisites for the modular organization of movements at the cortical level (Dubbioso, Raffin, Karabanov, Thielscher, & Siebner, 2017; Gentner & Classen, 2006).

However, despite the great potential of nTMS mapping, it is still underused for the assessment of motor plasticity, particularly in clinical settings. In order to use this approach in clinic, its reliability should be first established, both relative – allowing the stratifying of subjects/patients and absolute – allowing the interpretation of changes at the individual level (Atkinson & Nevill, 1998). Articles dedicated to TMS motor mapping reliability are mostly focused on the relative reliability (Cavaleri, Schabrun, & Chipchase, 2018; Forster, Limbart, Seifert, & Senft, 2014; Jonker et al., 2019; Kraus & Gharabaghi, 2016; Malcolm et al., 2006; McGregor et al., 2012; Ngomo, Leonard, Moffet, & Mercier, 2012; Pitkänen et al., 2017; Plowman-Prine, Triggs, Malcolm, & Rosenbek, 2008; Sankarasubramanian et al., 2015; Sinitsyn et al., 2019; Sollmann et al., 2013; van de Ruit, Perenboom, & Grey, 2015; Weiss et al., 2013; Wolf et al., 2004; Zdunczyk, Fleischmann, Schulz, Vajkoczy, & Picht, 2013), while absolute reliability has been investigated much less frequently (Jonker et al., 2019; Ngomo et al., 2012; Sankarasubramanian et al., 2015; van de Ruit et al., 2015). As for the multiple-muscle TMS mapping, to the best of our knowledge, there are fewer reports in the literature (Cavaleri et al., 2018; Forster et al., 2014; Plowman-Prine et al., 2008; Weiss et al., 2013), and the reliability of inter-muscle interactions have not yet been addressed. Furthermore, another limitation, considering TMS MCR reliability, is the lack of an agreement on which MCR parameters should actually be studied. Apart from the MCR parameters commonly used and recommended in the TMS clinical guidelines, such as areas, volumes, COGs and hotspots (Rossini et al., 2015) there is a range of more sophisticated parameters, such as MCR shape (Pitkänen et al., 2017), discrete peaks per MCR (Beaulieu, Flamand, et al., 2017; Cavaleri, Schabrun, & Chipchase, 2017; Schabrun, Hodges, Vicenzino, Jones, & Chipchase, 2015), MCR excitability profile (EP) (Novikov et al., 2018; Raffin et al., 2015) etc., but the validity of these parameters has not yet been fully established.

In this study we conducted a systematic investigation of both the absolute and relative reliability of three hand muscle MCRs and their interactions in a homogenous group of healthy male volunteers using a comprehensive (≥170 points) individual brain nTMS mapping. Additionally, we studied the somatotopy gradient among the investigated hand muscle MCRs. Moreover, to complement standard MCR metrics, we used overlaps between MCRs, and differences of MCR EPs with the idea that it would be possible to identify muscle-specific MCR features suitable for probing in longitudinal multiple-muscle TMS mapping studies.

## 2. Materials and methods

Data supporting the findings of this study are available from the corresponding author upon reasonable request.

### 2.1 Participants

22 young (19-33 y.o.) healthy male volunteers were enrolled in the study. Exclusion criteria were: any history of neurological/psychiatric disorders including fainting spells, any medications intake, cardiac implants, metallic implants in the head, implanted pumps, stimulators and shunts or MRI incompatibility for any reasons. We also strictly excluded subjects with special motor skills, for example professional or trainee sportsmen, musicians, surgeons, painters and people with special hobbies requiring high manual dexterity. Participants self-reported for the usual 6-9 hours of sleep before the TMS procedure, no alcohol intake 24 hours before, and the usual amount of coffee intake. Subjects self-reported their handedness. All subjects gave written informed consent in accordance with the Declaration of Helsinki. All subjects were screened for contraindications to TMS (Rossi, Hallett, Rossini, & Pascual-Leone, 2009) before the consent process. Experiments were approved by the IRB of the Research Center of Neurology N312 and the local Ethics Committee of HSE University, Moscow.

### 2.2 Overall experimental approach of test-retest nTMS mapping

Two nTMS mapping sessions were performed on each subject (day 1 and day 2) separated by 5-10 days. Both TMS mapping sessions were conducted for each volunteer at the same day time (morning/afternoon/evening). During the period between days 1 and 2 volunteers were asked to keep their lifestyle unchanged, in particular, not to change their usual motor activity (e.g., their usual fitness schedule), not to start learning new motor skills etc. Because of the violations of some of these requirements, we excluded four volunteers; therefore the data of 18 volunteers were used for further analysis.

### 2.3 Electromyography

MEPs were obtained from three upper limb muscles: two intrinsic hand muscles – abductor pollicis brevis (APB) and abductor digiti minimi (ADM) and one extrinsic hand muscle – extensor digitorum communis (EDC). Surface EMG was recorded by the integrated EMG device of the eXimia system (3 kHz sampling rate, band-pass filter of 10-500 Hz). Bipolar surface 0.6-cm^2^ ECG Ag-AgCl electrodes (3M Red Dot) were placed using belly-tendon montage: the active electrode was located over the muscle belly, the referent one-two cm distally, and the ground one – on the wrist of the same hand. APB and EDC muscles were chosen because they are usually associated with hand motor deficits, for instance, in stroke patients (thumb abduction and finger extension) (Nazarova et al., 2019), also APB is one of the most commonly investigated muscles in TMS studies (Rossini et al., 2015). ADM was added to trace the within-hand cortical somatotopy. The MEP peak-to-peak amplitudes were calculated online using eXimia software. During the initial preprocessing, EMG data were visually inspected; trials were rejected if noise or the preactivation of any muscle was higher than 20 µV (peak-to-peak EMG amplitude).

### 2.4 nTMS cortical mapping

All TMS mapping procedures were carried out in accordance with the TMS safety guidelines (Rossi et al., 2009). Prior to the mapping, individual anatomical T1-weighted magnetic resonance images were acquired by a 1.5T MR-scanner Siemens Magnetom Avanto (structural T1-weighted images, MPRAGE, 1 mm isotropic voxel, acquisition matrix 256×256). Single pulse nTMS investigation was performed using a figure-of-eight biphasic coil (outer winding diameter 70 mm) using a Nexstim nTMS device (Nexstim Ltd, Helsinki, Finland, version 3.2.2).

As a first step a “rough TMS mapping” (Krieg, 2017) procedure, starting from the “hand knob” region in the precentral gyrus (Yousry et al., 1997) was performed to find a “technical hotspot” for the APB muscle for further resting motor threshold (RMT) probing. The “technical hotspot” was defined as the coil position (considering the electric field (EF) direction) resulting in the highest MEP amplitudes from the APB muscle. Then RMT for the given “technical hotspot” was determined as a minimal stimulator output, producing contralateral APB MEPs with minimal amplitude of 50 µV in a resting muscle, in 5 out of 10 stimuli (Rossini et al., 2015). For the day 2 TMS mapping session we looked for the APB hotspot and RMT again.

The intensity of the stimulation during TMS mapping was 110% of the RMT for APB. The time lag between the stimuli varied randomly between 3 and 10 seconds. Each of the two mapping sessions included five sub-sessions, separated by 2-10 minutes. Each sub-session consisted of 49-79 TMS pulses (a constant number of points for each volunteer). The stimulation nodes were pre-set using a virtual MRI-based grid in the Nexstim navigation software with 5×5 mm2 squares. We kept the coil perpendicular to the closest segment of the sulcus (Bashir, Perez, Horvath, & Pascual-Leone, 2013; Krieg et al., 2017; Raffin, Pellegrino, Di Lazzaro, Thielscher, & Siebner, 2015).

The outer margin of a given MCR was determined if no MEP could be elicited in two consecutive stimulation trials. During TMS mapping sub-sessions, the same points were stimulated in the forward and reverse order (forward-reverse-forward-reverse-forward) to decrease the effect of the regular stimulation of the same point (Figure 1). On day 2, the points were stimulated in exactly the same order as on day 1. An error of the navigation for each cortical spot was kept below 2 mm and the tilt was constant, according to the navigation system feedback. In the end, the whole stimulation session consisted of 177-395 stimuli per day depending on the volunteer (see an example of APB MCR in Figure 1).

**FIGURE 1.**
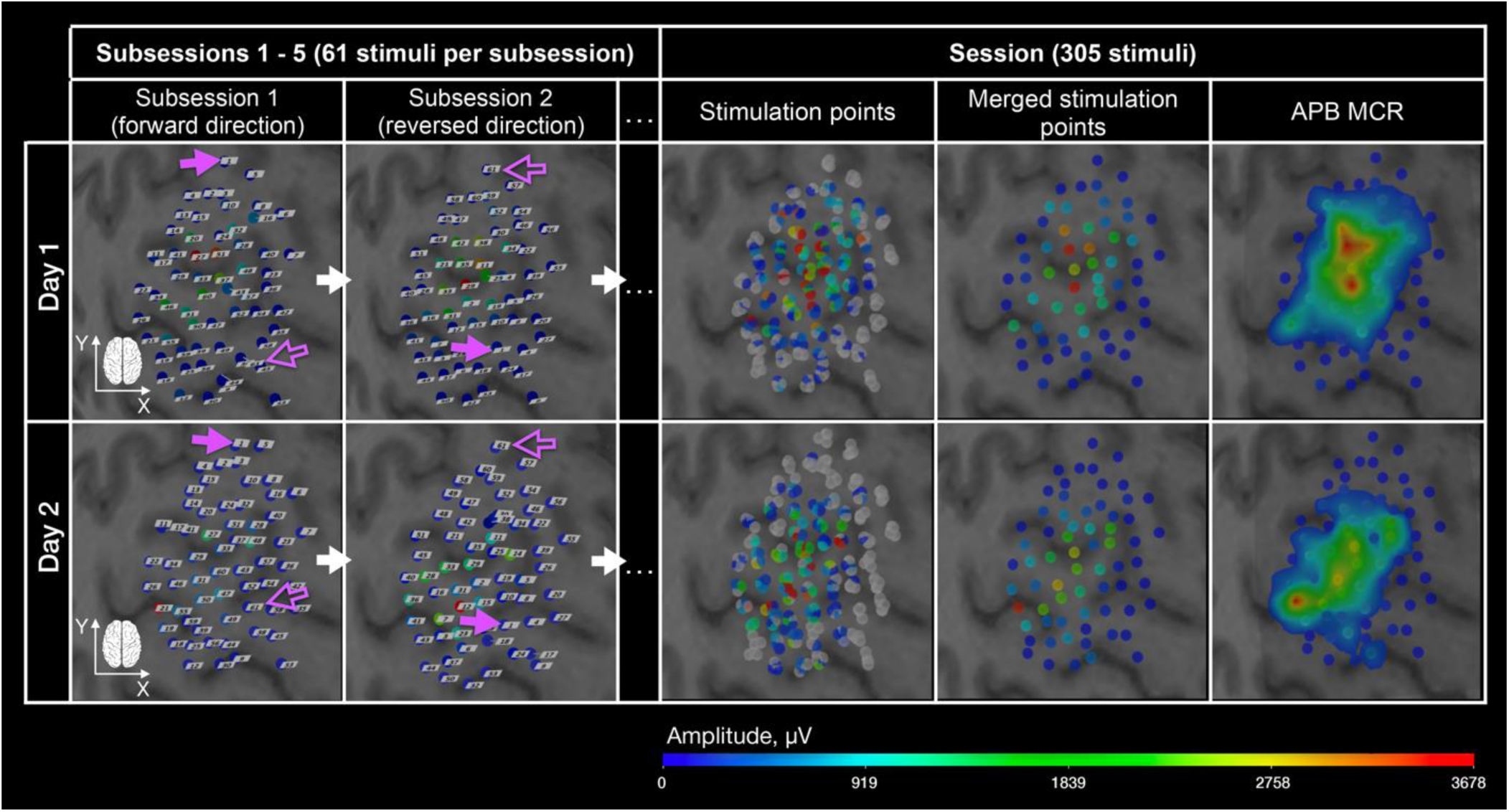
nTMS of the left primary motor cortex for a representative subject (visualization in TMSmap software). Stimulation points are shown for both days (upper raw – day 1, bottom raw – day 2). In the “Sub-sessions 1-5” column the order of the stimulation is shown. Magenta filled arrow indicates the first stimulation point, magenta non-filled arrow – the last stimulation point in a TMS mapping sub-session. One can notice that the location of the stimulation points for day 1 and day 2 are almost identical. “Stimulation points” column represents all stimulation points per day 1 and 2, while in the “Merged stimulation points” column, points after spatial filtering are shown. X-axis corresponds to the lateral to medial direction; Y-axis corresponds to the posterior to anterior direction. The color scale represents MEP amplitude in microvolts (µV).

#### 2.4.1 Along central sulcus nTMS mapping

Considering that the coil in our study was kept perpendicular to the central sulcus, we also aimed at comparing our results with those, obtained using the so-called “along-sulcus” TMS mapping (Raffin et al., 2015; Raffin & Siebner, 2019), although it was not our original intention. As using this “along-sulcus” TMS mapping approach, a clear within-limb somatotopy gradient between the first dorsal interosseus and ADM MCRs, was recently reported (Raffin et al., 2015; Raffin & Siebner, 2019). We manually identified stimulation points located in the “hand knob” along the central sulcus for day 1. We refer to these data as “along-sulcus” mapping, while for mapping data based on all points as “whole” MCRs.

#### 2.4.2 Single-muscle MCR size and topography parameter calculation

MCRs were constructed and analyzed using TMSmap – a freely available software for the quantitative assessment of TMS cortical mapping data, which we developed and described previously (Novikov, Nazarova, & Nikulin, 2018). We calculated commonly used size MCR parameters such as areas, volumes, mean MEP per MCR, and topography parameters such as hotspot and COG locations (the COG formula was the same as elsewhere, including the TMSmap software description (Novikov et al., 2018). We merged stimulation points located closer than 2 mm (see Figure 1) (Novikov et al., 2018) and used mean MEP amplitudes after merging for MCR construction. The threshold MEP amplitude for MCR construction was 50 µV for a merged point. For the purpose of MCR comparison we redefined the hotspot: instead of using the “technical hotspot” where RMT was defined, we considered the whole TMS mapping data per day. Thus, we again defined the “final hotspot” as the location over which averaged MEPs with the highest peak-to-peak amplitude were evoked in each of the target muscles, so like that we determined hotspots for all three muscles. Additionally, we compared the excitability profiles (EPs) of different MCRs using EMD metrics (see the next subsection).

#### 2.4.3 The topography parameters of multiple muscle MCR interactions

As a reflection of multiple muscle MCR interactions we used:

- Geodesic shifts between different muscles MCR COGs’ and hotspots’ x and y coordinates (reflecting shifts in the medio-lateral and anterior-posterior directions, respectively);
- Area and volume overlap between two MCRs. We used an approach for MCR overlap calculation implemented in TMSmap (Novikov et al., 2018). Briefly, area and volume normalized overlaps between MCRs were calculated using the following formulas:

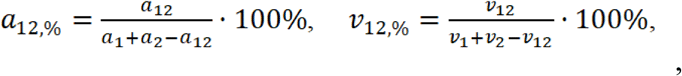

where a_1_, a_2_ are the areas of MCR 1 and 2 respectively; v_1_, v_2_ are the volumes of MCR 1 and 2 respectively; a_12_, v_12_ are the common area and volume for the MCR 1 and 2 respectively. So, the normalized overlaps between MCRs can be interpreted in such a way that the value of 100% indicates completely identical MCRs, while the value of 0% indicates no overlap between MCRs. Further in the text we will use only normalized overlaps and refer to them as overlaps.
- EPs of two MCRs of different muscles on one day were compared using the earth mover’s distance (EMD), which is also called the Wasserstein metric (Rubner, Tomasi, & Guibas, 1998). It characterizes a minimum cost of turning one distribution into the other (Rubner et al., 1998). It was calculated in TMSmap software as described previously (Novikov et al., 2018). The same approach of EMD calculation was used for the reliability assessment of MCR EPs between days.

### 2.4 MNI MCR construction

Additionally to individual MRI-based MCR analysis, we probed the within-limb somatotopy of the MNI normalized MCRs. For this we transformed MCRs to MNI space, using a custom-made algorithm utilizing the SPM8 software (The Wellcome Department of Imaging Neuroscience, Institute of Neurology, University College London, UK) and TMSmap (Novikov et al., 2018).We used only 17 right-handed subjects (excluding one left-handed subject). After MNI normalization, we calculated the MNI MCR parameters in the same way, as we did for the individual space MCRs. In TMSmap we also constructed averaged MNI MCRs, averaging the MCRs of APB, ADM, and EDC across-subjects for both days. For this we normalized each MCR in MNI space by the maximal MEP amplitude for this MCR and obtained new MCR as a percentage of the individual MEP amplitude maximum, then these values were averaged across all the participants for each muscle for both days (Figure 6).

### 2.5 Statistical analysis

Apart from the analysis in TMSmap, further statistical analysis was performed using the SPSS package for Windows (IBM) and Matlab (Natick, USA). The level of significance was defined as α = 0.05. Family-wise error rate (FWER) correction was performed to control for multiple comparisons.

#### 2.5.1 MCR within-limb somatotopy gradient assessment

We used one-way repeated measures ANOVA (rmANOVA) to confirm the within-limb gradient among hand muscle MCRs (for this we again used 17 right-handed subjects, excluding one left-handed participant). We compared COGs of the different muscle MCRs to probe the mediolateral shift among them for (1) the individual “whole” MCRs, (2) the “along-sulcus” mapping data and (3) the MNI version of MCRs. Additionally, using two-way rmANOVA we compared geodesic shifts between different muscle MCR COGs inside one day for the “whole” and the “along-sulcus” mapping approaches (factors: muscle, TMS mapping approach). Finally, we used one-way rmANOVA to compare the areas of different muscle MNI MCRs for each day. Shapiro-Wilk test for normality (Ghasemi & Zahediasl, 2012) and Mauchly’s test for sphericity assessment (Beddo & Kreuter, 2004) were applied.

#### 2.5.2 Reliability assessment

Before the reliability analyses, we also assessed normality by Shapiro-Wilk test (Ghasemi & Zahediasl, 2012) and homoscedasticity, using Levene’s test (Beddo & Kreuter, 2004; Weir, 2005). When non-normality or heteroscedasticity were found for more than one muscle, a natural logarithmic transformation was applied before reliability analysis for this parameter for all three muscles (Atkinson & Nevill, 1998; Beaulieu, Flamand, Massé-Alarie, & Schneider, 2017). In case when after logarithmic transformation the normality still was not achieved we decided not to do bootstrapping, considering that in case of low amount of data it is difficult to estimate the distribution’s tails (Orloff & Bloom, 2014), and in this case we did not estimate SDC. The general pipeline of MCR reliability assessment is presented in Figure 2.

**FIGURE 2.**
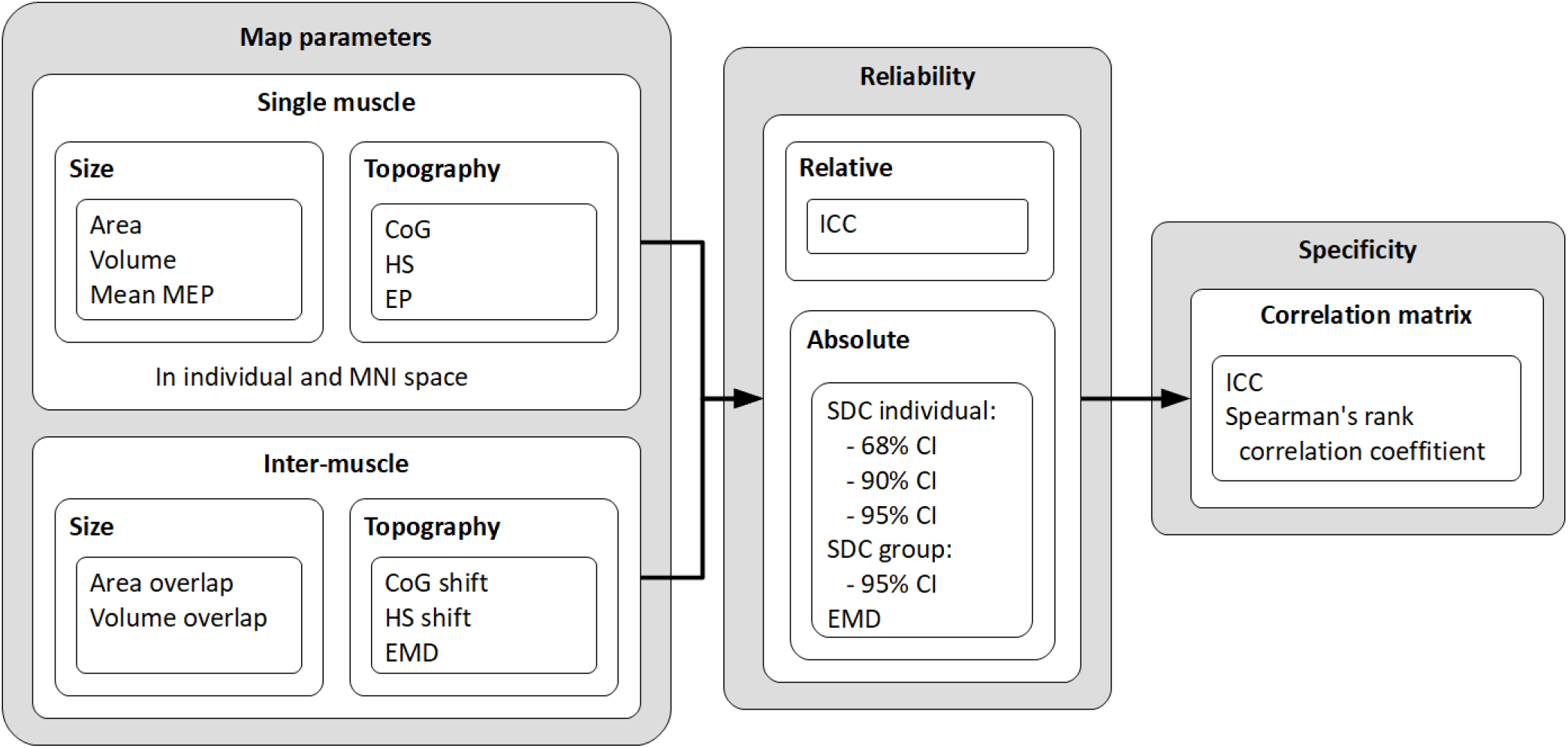
A general pipeline of the reliability assessment.

#### 2.5.3. Relative reliability assessment (ICC)

We used a two-way mixed model absolute agreement intraclass correlation coefficient (ICC) to assess relative reliability for single-muscle MCR parameters (Beaulieu, Flamand, et al., 2017; Koo & Li, 2016). We calculated ICC between days for RMTs, map areas, volumes, mean MEP, hotspots” and COGs” x and y coordinates for each MCR. To check the reliability of multiple muscle MCR relationship we calculated (1) the shifts between different muscles MCRs’ COGs’ and hotspots’ coordinates, (2) the extents of the different muscle MCR overlaps and (3) EMDs between the different muscle MCR EPs. For relative reliability description we used the following ranges: below 0.50 – poor; between 0.50 and 0.75 – moderate; between 0.75 and 0.90 – good; above 0.90 – excellent (Koo and Li 2015).

#### 2.5.4 Muscle-specificity assessment

In addition to relative reliability, we investigated whether MCR parameters were muscle-specific. For this, in addition to ICC, we assessed the Spearman rank correlation for the same MCR parameters. We considered a parameter to be muscle-specific when its values for a given muscle between days, agreed (ICC) or correlated (Spearman correlation coefficient) significantly more for the same-muscle MCR, compared to when this parameter was evaluated for the MCRs of other muscles (either on the same day or between days). In other words, muscle-specificity provides the possibility of identifying the correct muscle, among other muscles, using a certain MCR parameter.

#### 2.5.5. MCR EP comparison using EMD

To investigate the validity of different MCR EPs we compared:

(1) normalized EMD for the same muscle MCRs between days, calculated according to the formula (an example for APB MCR):

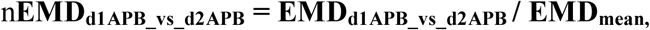

where

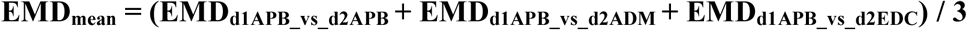

and (2) normalized EMD across-muscle MCRs between days, calculated according to the formula (an example for APB MCR versus two other muscles’ MCRs):

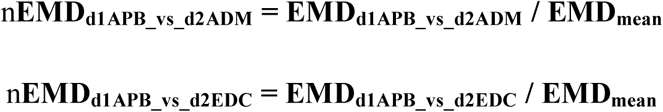

To compare normalized EMDs between same-muscle MCR EPs across days versus different muscle MCR EPs across days we used the non-parametric Mann-Whitney U test as the distribution of the residuals in one of the samples was not normal according to Shapiro-Wilk test.

#### 2.5.6. Absolute reliability (SDC) assessment

For the assessment of the absolute reliability of the MCR parameters we used the smallest detectable change (SDC, a.k.a. minimal detectable change) calculation. SDC represents the minimal change that a subject must show on the scale to ensure that the observed change is real and not just a measurement error (Beaulieu, Flamand, et al., 2017; Weir, 2005). We estimated SDC using the standard error of measurements (SEMeas):

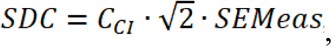

where *C*_*CI*_ is a coefficient which is equal to 1.00, 1.65 or 1.96 associated with a 68%, 90% and 95% confidence intervals (CI), respectively. The 68% CI was included because we assume that it may be adequate for change interpretation at the individual level. 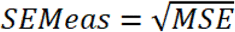, where MSE is the mean squared error obtained from the rmANOVA applied on test and retest measurements (Weir, 2005). When SDC values were obtained for log-transformed parameters, we then performed the antilog transformation as in (Beaulieu, Massé-Alarie, Ribot-Ciscar, & Schneider, 2017).

## Results

Figure 3 provides an example of a representative subject’s test-retest single-muscle MCRs and their overlaps for both days. The figure shows the distribution of MEP amplitudes depending on the stimulation location relative to the mapping grid. One can clearly see that all MCRs are different but it is also hard to categorize this difference visually. In this example, the APB MCR is shifted laterally compared to two other MCRs (COG x is smaller), it also has the biggest size; and the overlap between APB and ADM MCRs is the most prominent. We provide the data for all the participants’ single-muscle MCR parameters from TMSmap software (areas, volumes, mean MEPs, hotspot and COG location) and the inter-muscle MCR interaction parameters (areas and volumes overlaps and EMDs between the different muscle MCRs) (Tables S7-S9 in the Supporting material).

**FIGURE 3.**
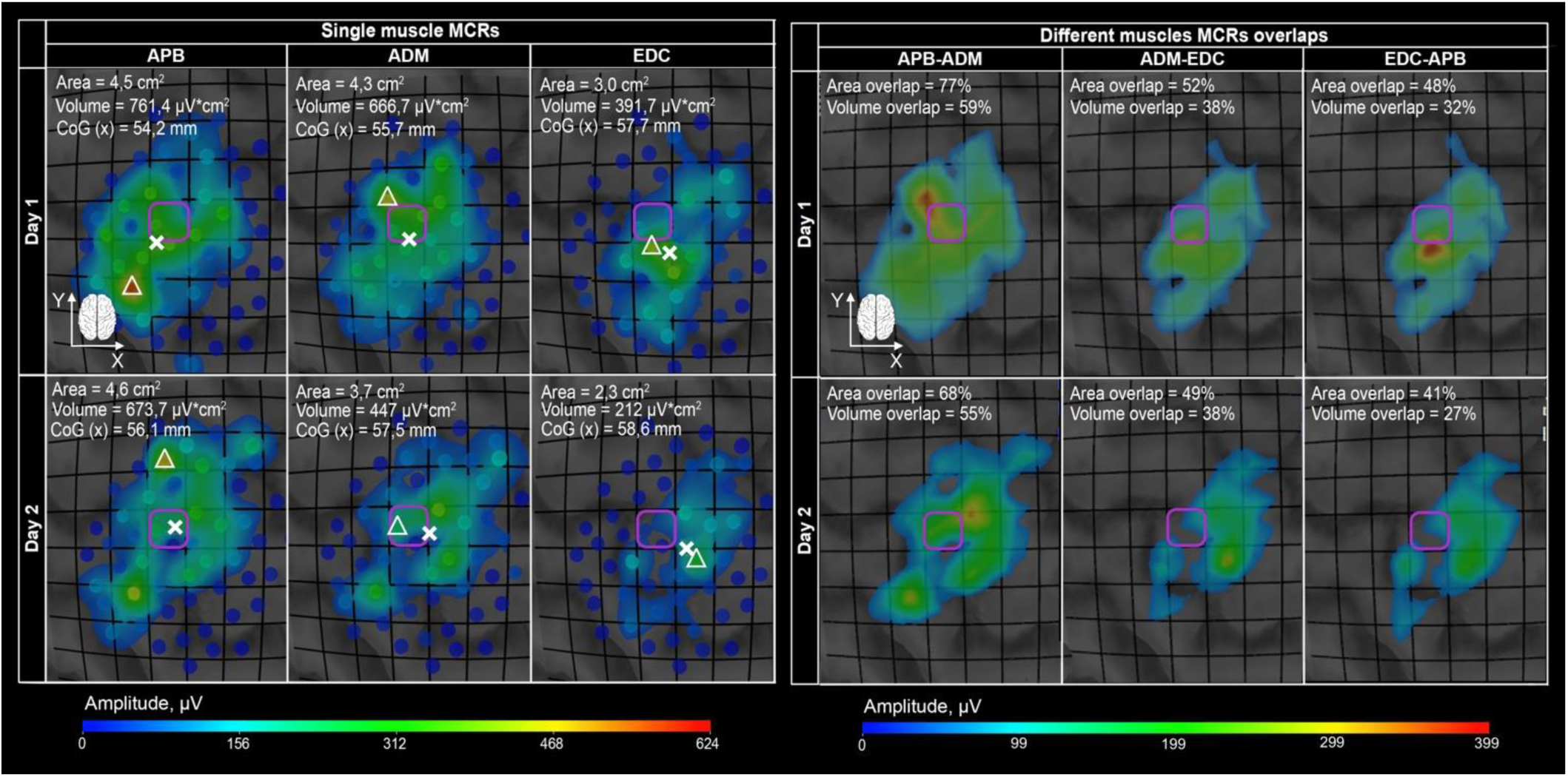
Exemplary TMS MCRs of APB, ADM and EDC muscles for both days in a representative subject. MCR reconstructions are based on 305 points per day; points located closer than 2 mm are merged. The areas and volumes and areas and volumes overlaps (in %) for MCR pairs are shown. Magenta squares indicate the same cortical area for all MCRs. COGs are shown with white crosses, hotspots – with white triangles. X-axis corresponds to the lateral to medial direction; Y-axis corresponds to the posterior to anterior direction. The color scale represents MEP amplitude in microvolts (µV).

The results are described in the following manner. First, we report the within-limb somatotopy gradient for the individual MRI MCRs and then for MNI data. Second, we show relative and absolute reliability and the muscle-specificity of different MCR parameters, assessed for all single-muscle MCRs. Finally, we present the inter-muscle MCR interaction reliability in the same order as for a single-muscle MCRs (see the general pipeline of the reliability assessment in Figure 2 in Methods section).

### 3.1. Within-limb somatotopy gradient

#### 3.1.1. Somatotopy gradient for individual MRI data

Despite the great overlap between the muscle MCRs, we observed a mediolateral shift between the APB MCR and the two other muscle MCRs, based on COGs’ x coordinates. This shift was significant for day 1 (1 way rmANOVA: day 1: p < 10^−6^ (pair-wise comparison after FWER APB-ADM: p < 10^−5^, ADM-EDC: p = 0.227, APB-EDC: p < 10^−3^). For the second day this shift was observed only as a tendency (p = 0.021 (pair-wise comparison after FWER APB-ADM: p = 0.064, ADM-EDC: p = 0.539, APB-EDC: p =0.071)).

#### 3.1.2. Along-sulcus TMS mapping

We compared our results with those obtained using the “along-sulcus” mapping (Raffin et al., 2015; Raffin & Siebner, 2019) because previously for this approach within-limb somatotopy gradient between the first dorsal interosseus and ADM muscle MCRs was reported (Raffin et al., 2015; Raffin & Siebner, 2019). We were able to identify 28 to 63 points per participant in the precentral gyrus along the central sulcus (see an example in Figure 4). In two participants all three muscle MCRs were shifted rostrally beyond the central gyrus, and, thus, there were no MEPs ≥ 50 µV in the points along the central sulcus (see an example in Figure 5). When comparing the mediolateral shift of the “along-sulcus” MCRs, based on day 1 COG x-coordinates in 15 participants with no null along-sulcus MCRs, we observed a significant difference between the APB MCRs versus the ADM and EDC MCRs COG x-coordinates (1-way rmANOVA p < 10^−4^, pair-wise comparison after FWER – APB-ADM: p = 0.001, ADM-EDC: p = 1, APB-EDC: p < 10^−3^). To check whether the mediolateral somatotopy gradient was more or less pronounced in the “along-sulcus” in contrast to the “whole” MCRs, we compared the differences between the COG x-coordinates between the different muscle MCRs for these two mapping approaches. We found that the mediolateral somatotopy gradient was greater for the “whole” MCRs compared to the “along-sulcus” MCRs (rmANOVA with two factors (“muscle-combination” and “TMS mapping type”) showed a significant factors interaction (p = 0.005). While looking at the example in Figure 4, one can see that the size of different muscle MCRs for “whole” and the “along-sulcus” mapping does not correspond. While the “whole” APB MCR is bigger than the “whole” ADM MCR, for the “along-sulcus” MCRs this interrelation is opposite. Considering (1) this mismatch, (2) the fact that it was impossible to obtain ≥70 stimulation points along the central sulcus (in analogy with 10 repetitions of 7 targets as in (Raffin & Siebner, 2019)) in any participant and (3) that in some participants we obtained no MEPs along the central sulcus at all, we have not further analyzed the “along-sulcus” MCRs quantitatively.

**FIGURE 4.**
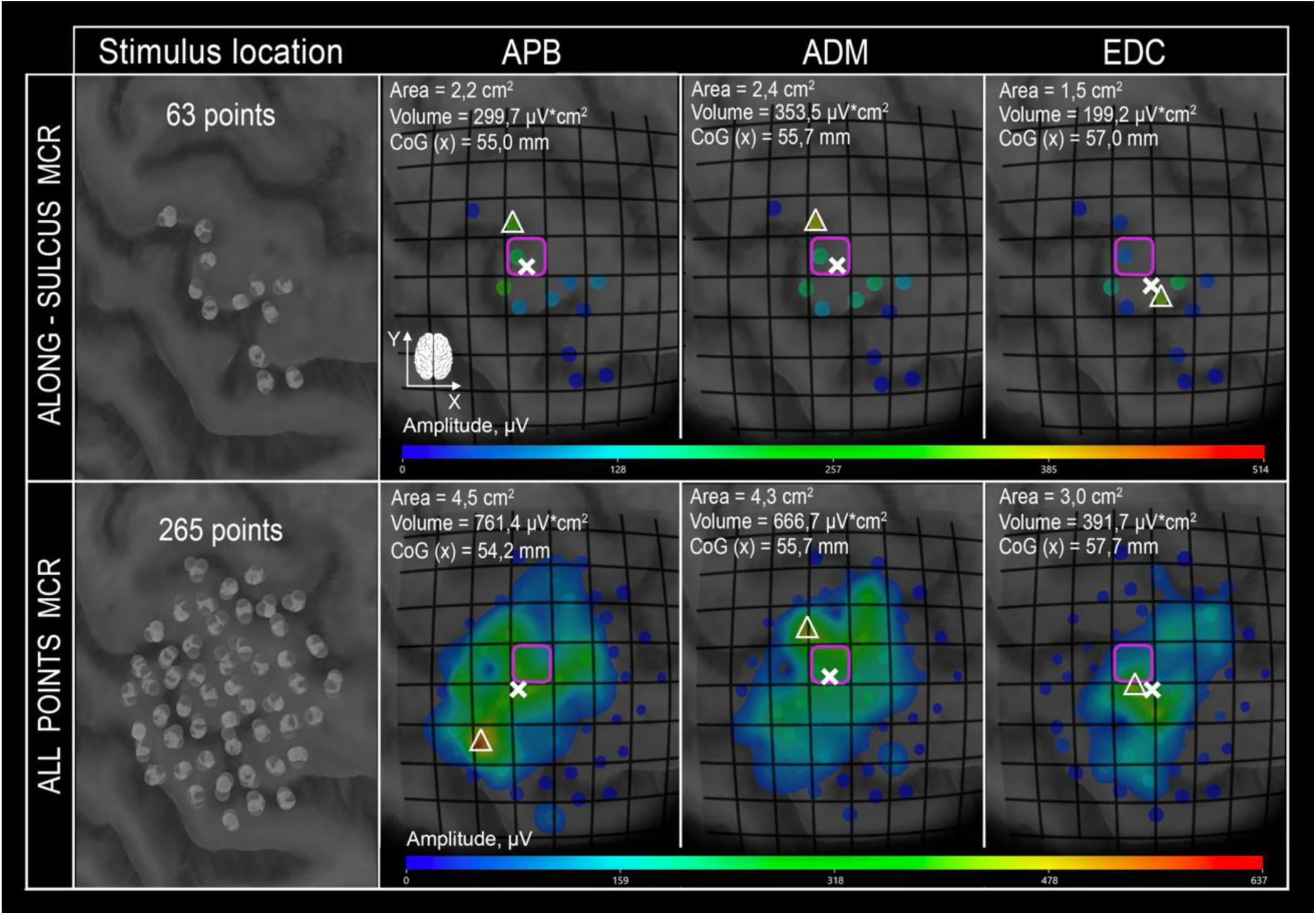
Comparison between the “whole” and the “along-sulcus” TMS mapping data in a representative subject for day 1. In the upper row “along-sulcus” mapping results for APB, ADM and EDC muscles are shown. The lower row represents the “whole” MCRs for all three muscles. COGs are shown with white crosses, hotspots – with white triangles. One can notice that in the “whole” TMS mapping case the APB MCR area is bigger than ADM MCR area, but in the “along-sulcus” mapping case this interrelation is opposite. Magenta squares indicate the same cortical area. X-axis corresponds to the lateral to medial direction; Y-axis corresponds to the posterior to anterior direction. The color scale represents MEP amplitude in microvolts (µV).

**FIGURE 5.**
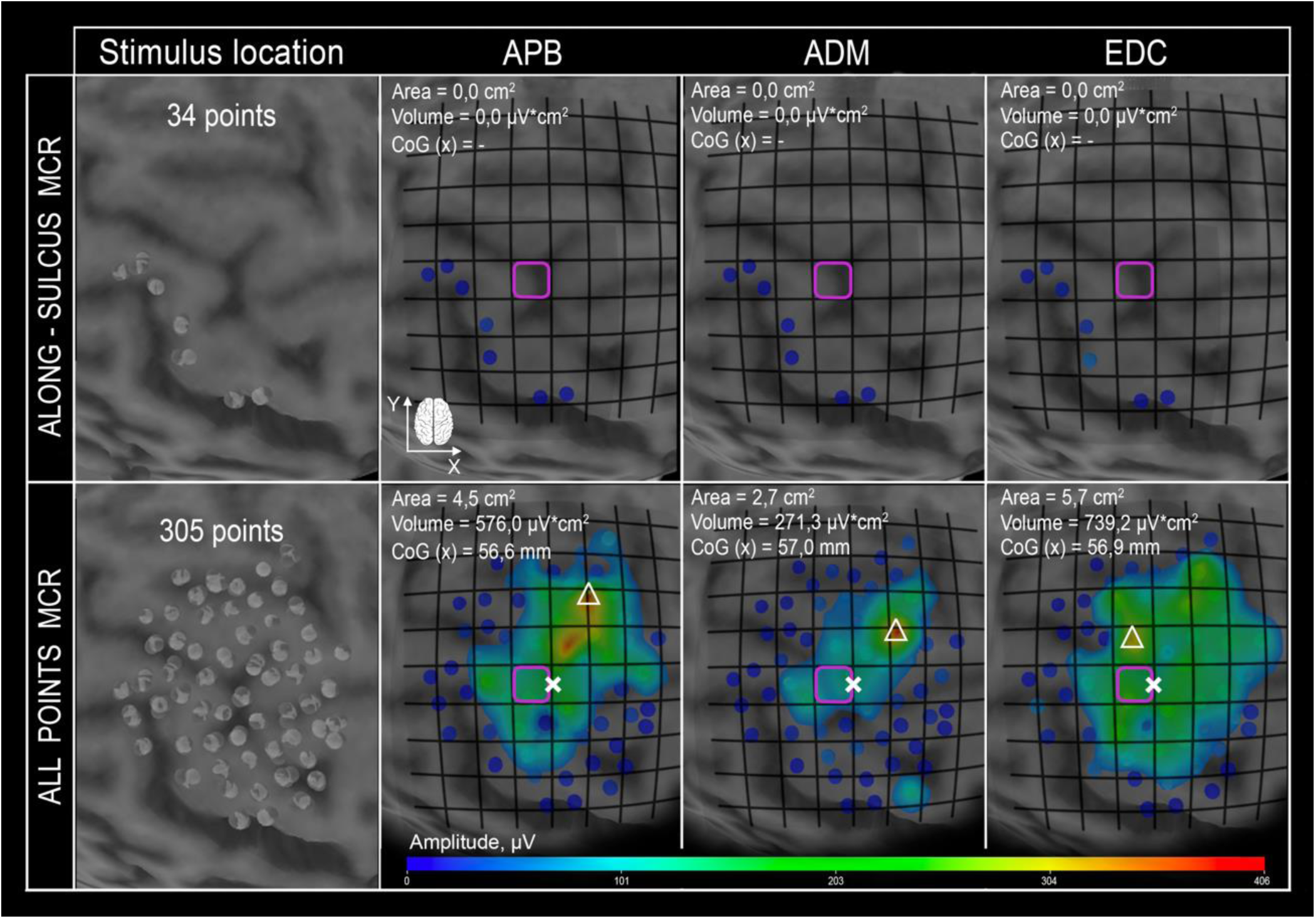
An example of TMS mapping dataset where there are no MEPs > 50µV along the central sulcus. In the upper row along-sulcus mapping data for all three muscles are shown, no MEP with amplitude >50µV can be seen. The lower row represents the “whole” MCRs for all three muscles. COGs are shown with white crosses, hotspots – with white triangles. The color scale represents MEP amplitude in microvolts. Magenta squares indicate the same cortical area. X-axis corresponds to the lateral to medial direction; Y-axis corresponds to the posterior to anterior direction.

**FIGURE 6.**
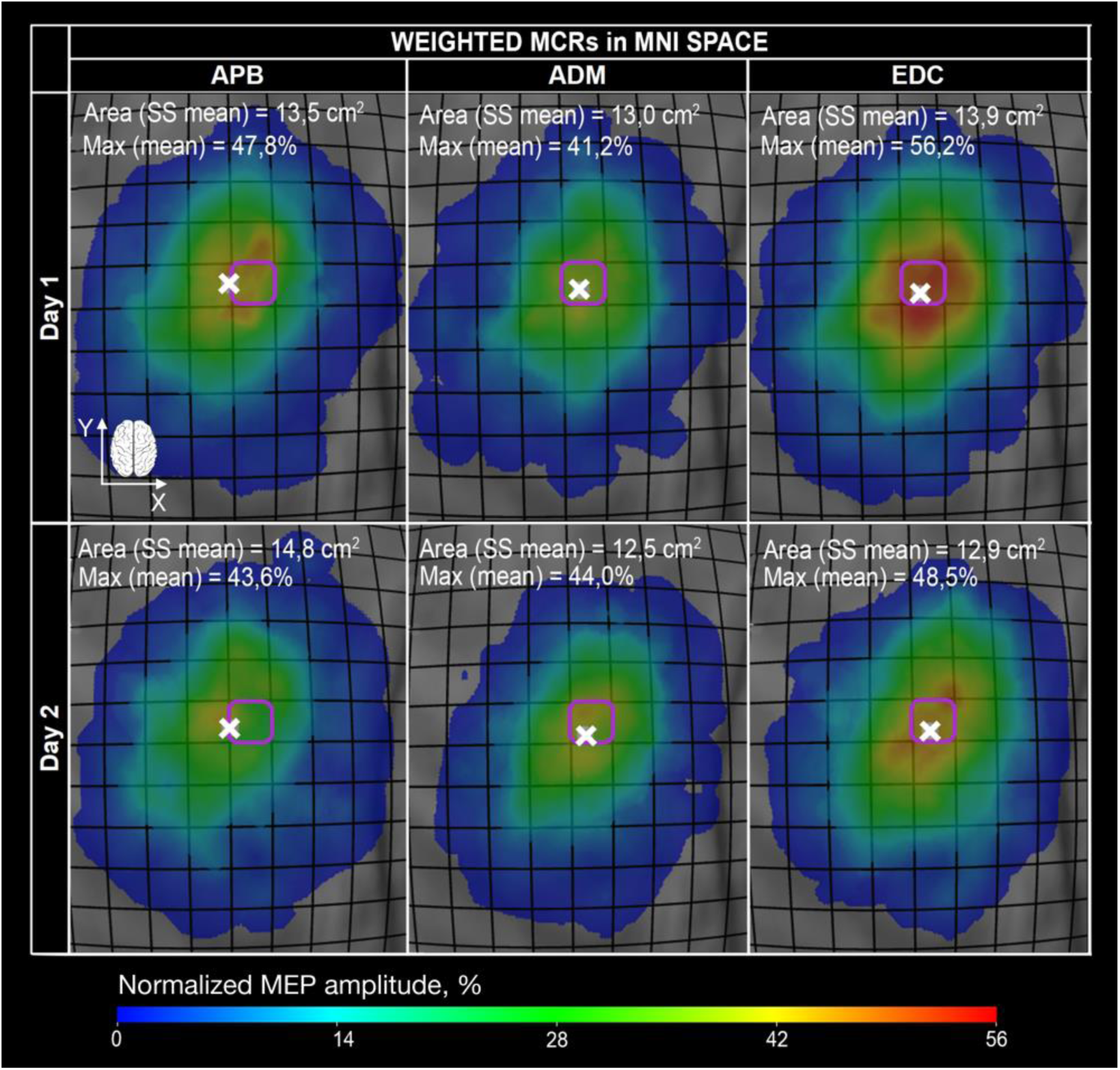
Weighted MNI MCRs of APB, ADM and EDC, day 1. COGs are shown with white crosses. Weighted MCR areas (in cm^2^) and maximum of the overlapped MEPs (in %) are shown. Magenta squares indicate the same cortical area. X-axis corresponds to the lateral to medial direction; Y-axis corresponds to the posterior to anterior direction. The color scale reflects the mean weighted MEP at a stimulation point.

#### 3.1.3. MNI data somatotopy gradient

In addition to individual data assessment, we also probed the somatotopy gradient of MCRs across subjects, using data co-registered to the MNI space. For MNI assessment we again used 17 right-handed subjects. We observed that APB MCR were more lateral compared to the ADM and EDC MCR for both days based on COG x-coordinates (rmANOVA after FWER APB-ADM day 1: p = 0.001, day 2: p = 0.016, APB-EDC day1: p < 10^−3^, day2: p = 0.004). We also found the difference between ADM and EDC MCR areas: ADM MCR area was significantly smaller for both days (1-way rmANOVA, pair-wise comparison after FWER, day1: p=0.005, day2: p=0.041).

In addition, we created weighted normalized MCRs for both days (Figure 6). Normalized MEPs amplitudes (averaged across subjects) varied from 0 to 48%, meaning that not more than half of the subjects had their maximum MEPs in the same spot. Consistently with the previously observed somatotopy gradient, we again found a lateral shift of the COG of the APB MCR compared to ADM and EDC MCRs. It can also be seen that EDC normalized MCR has a bigger red zone, indicating that the optimal areas with high MEPs for EDC MCRs across subjects are located closer to each other compared to the two other muscle MCRs.

### 3.2. Single-muscle cortical TMS map reliability analysis

#### 3.2.1. Relative reliability

##### 3.2.1.1. TMS map size parameters

ICC for areas and log-transformed volumes and mean MEPs varied from good to excellent (0.71 - 0.85), depending on a muscle (Table 1).

**TABLE 1.** Relative (ICC) and absolute (SDC) reliability values for single-muscle (APB, ADM and EDC) MCR size parameters in the individual space. The data are shown for different confidence intervals (CI)

| Parameter, muscle | ICC | SEMeas | SDC |  |  |  |
| --- | --- | --- | --- | --- | --- | --- |
|  |  |  | Individual |  |  | Group<br>(n=18) |
|  |  |  | 68% CI | 90% CI | 95% CI | 95% CI |
| RMT % |  |  |  |  |  |  |
| APB | 0.99 | 0.48 | 0.7 | 1.1 | 1.3 | 0.3 |
| Area, cm<sup>2</sup> |  |  |  |  |  |  |
| APB | 0.85 | 0.72 | 1.03 | 1.69 | 2.01 | 0.47 |
| ADM | 0.84 | 0.76 | 1.08 | 1.78 | 2.11 | 0.50 |
| EDC | 0.70 | 0.93 | 1.32 | 2.17 | 2.58 | 0.61 |
| Ln (Mean MEP (mp)), μV | | | | | | |
| APB | 0.72 | 0.296<br>($\times/\div 1.34$) | 0.419<br>($\times/\div 1.52$) | 0.691<br>($\times/\div 2.00$) | 0.821<br>($\times/\div 2.27$) | 0.193<br>($\times/\div 1.21$) |
| ADM | 0.79 | - | - | - | - | - |
| EDC | 0.74 | 0.293<br>($\times/\div 1.34$) | 0.414<br>($\times/\div 1.51$) | 0.684<br>($\times/\div 1.98$) | 0.812<br>($\times/\div 2.25$) | 0.191<br>($\times/\div 1.21$) |
| Ln (Volume) |  |  |  |  |  |  |
| APB | 0.79 | 0.497<br>($\times/\div 1.64$) | 0.703<br>($\times/\div 2.02$) | 1.160<br>($\times/\div 3.19$) | 1.377<br>($\times/\div 3.96$) | 0.325<br>($\times/\div 1.38$) |
| ADM | 0.85 | - | - | - | - | - |
| EDC | 0.72 | 0.425<br>($\times/\div 1.53$) | 0.602<br>($\times/\div 1.82$) | 0.993<br>($\times/\div 2.70$) | 1.179<br>($\times/\div 3.25$) | 0.278<br>($\times/\div 1.32$) |

##### 3.2.1.2 TMS map standard topography parameters (hotspots, COGs)

ICC for hotspots and COGs was almost always excellent for individual MRI data (>=0.9) (Table 2). In the MNI space ICC decreased: for COGs it varied from good to excellent, while for hotspots - from fair to good (Table 2). We suggest that MNI ICC for hotspots and COGs is a more correct because it is not sensitive to high data variance among the individual brains.

**TABLE 2.** Relative (ICC) and absolute (SDC) reliability values for hotspots and COGs for single-muscle (APB, ADM and EDC) MCRs in the individual space for different confidence intervals (CI)

| Parameter, muscle | ICC | SEMeas | SDC |  |  |  |
| --- | --- | --- | --- | --- | --- | --- |
|  |  |  | Individual |  |  | Group<br>(n=18) |
|  |  |  | 68% CI | 90% CI | 95% CI | 95% CI |
| COG x, mm |  |  |  |  |  |  |
| APB | 0.99 | 1.79 | 2.53 | 4.17 | 4.96 | 1.17 |
| ADM | 0.98 | 2.28 | 3.23 | 5.33 | 6.33 | 1.49 |
| EDC | 0.99 | 1.54 | 2.17 | 3.58 | 4.26 | 1.00 |

| COG y, mm |  |  |  |  |  |  |
| --- | --- | --- | --- | --- | --- | --- |
| APB | 0.96 | 1.72 | 2.44 | 4.02 | 4.77 | 1.13 |
| ADM | 0.95 | 1.95 | 2.76 | 4.55 | 5.41 | 1.27 |
| EDC | 0.95 | 1.86 | 2.64 | 4.35 | 5.17 | 1.22 |
| hotspot x, mm |  |  |  |  |  |  |
| APB | 0.96 | 3.57 | 5.05 | 8.33 | 9.89 | 2.33 |
| ADM | 0.97 | 3.06 | 4.33 | 7.15 | 8.49 | 2.00 |
| EDC | 0.97 | 2.81 | 3.97 | 6.56 | 7.79 | 1.84 |
| hotspot y, mm |  |  |  |  |  |  |
| APB | 0.74 | 4.41 | 6.23 | 10.29 | 12.2 | 2.88 |
| ADM | 0.86 | 3.03 | 4.29 | 7.08 | 8.41 | 1.98 |
| EDC | 0.73 | 4.88 | 6.90 | 11.38 | 13.5 | 3.19 |

#### 3.2.2. Absolute reliability

##### 3.2.2.1. MCR size parameters

Individual SDC (CI 95%) for areas varied from 2.01 to 2.58 cm^2^. When the CI is decreased to 68%, individual SDC for areas was half the size: 1.03-1.32 cm^2^, depending on the muscle. Evidently, a similar dependency of the SDCs on CI has been also observed for all other parameters (Table 1).

##### 3.2.2.2. MCR standard topography parameters (COGs and hotspots)

Individual SDC (CI 95%) for the hotspot coordinates was around 1 cm, while for COG coordinates – around 0.5 cm. SDC values for different muscle MCRs were very similar (Table 2).

##### 3.2.2.3. MCR excitability profiles comparison (EMD)

Additionally, we assessed the reproducibility of a given muscle MCR EP, using EMD (Rubner et al., 1998), implemented in the TMSmap software (Novikov et al., 2018). To our knowledge, this is the first time this metric was used for a quantitative comparison of the TMS MCRs in a test-retest study. We showed that the between-days EMD is significantly smaller for the same muscle MCR EPs (thus indicating higher reproducibility) than for the different muscle MCR EPs (Mann-Whitney U test, p < 10^− 4^).

#### 3.2.3. Muscle-specificity

##### 3.2.3.1. MCR size parameters

Based on ICC, none of the MCR size parameters was muscle-specific (Figure 7). When using Spearman’s correlation, however, we observed muscle specificity after FWER correction - this can be seen as stronger correlation values on the main diagonal of the matrix compared to off-diagonal values, reflecting a correlation between different muscles’ MCR parameters values (Figure 7). The situation for the MNI data was similar, mean MEP muscle-specificity was significant (Figure 8).

**FIGURE 7.**
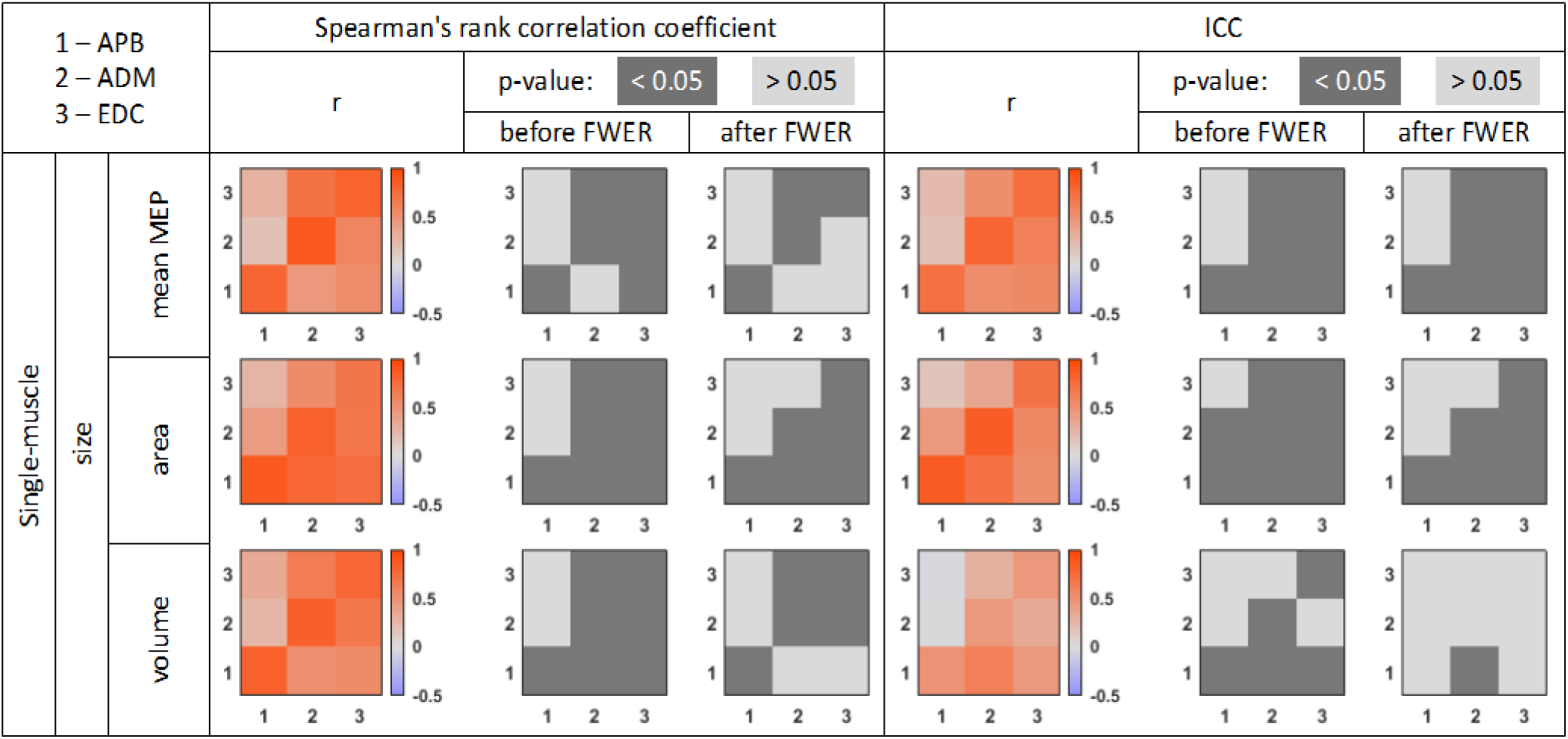
Correlation matrices for the size parameters of a single-muscle MCRs in the individual space (1 – APB, 2 – ADM, 3 – EDC). Diagonal elements in the matrix indicate within-muscle comparison, while off-diagonal elements indicate between-muscle comparisons.

**FIGURE 8.**
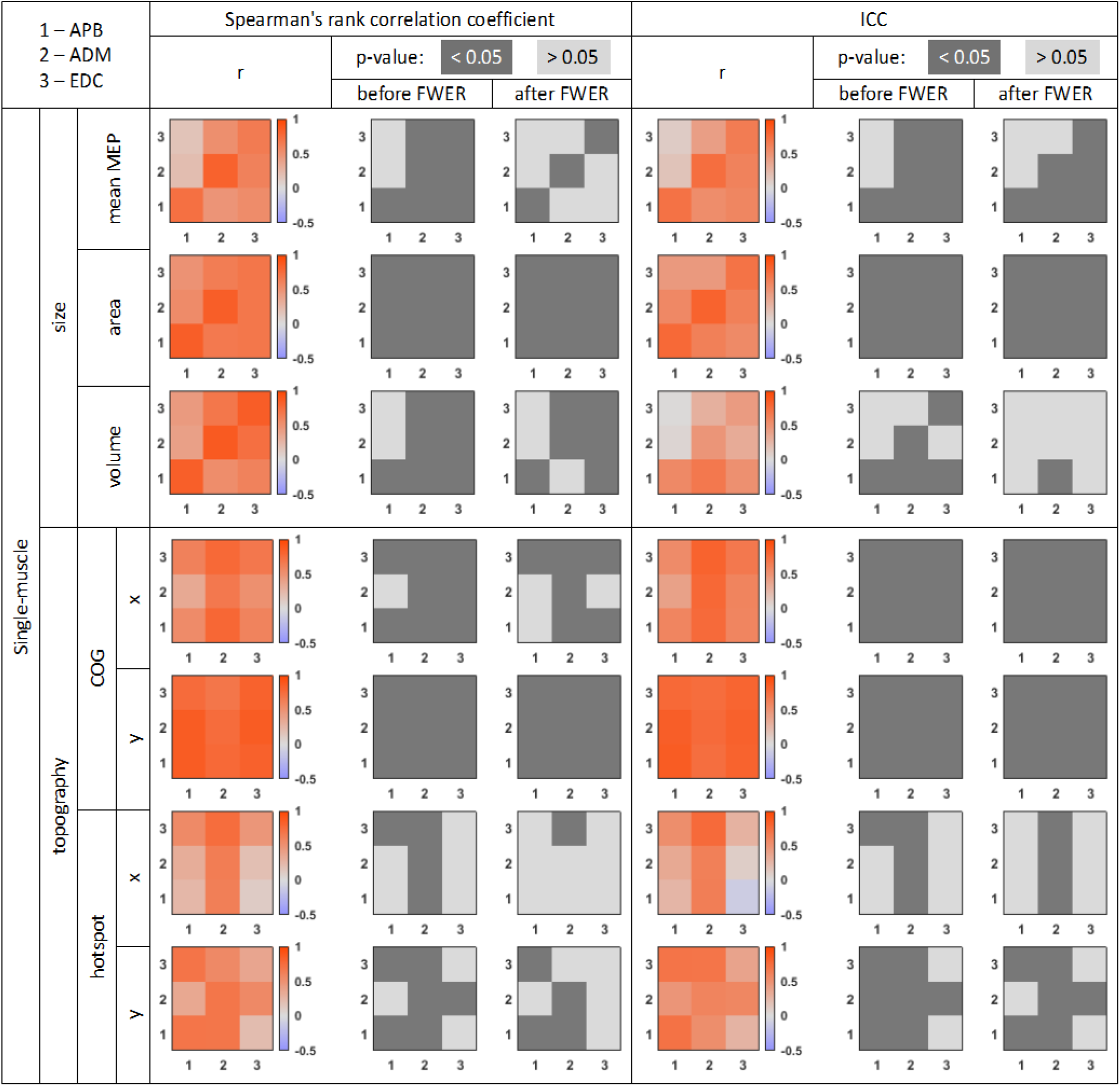
Correlation matrices for topography parameters of single-muscle MCRs in the MNI space (1 – APB, 2 – ADM, 3 – EDC). The color bar indicates the strength of the correlation. Diagonal elements in the matrix indicate within-muscle MCR comparison, while off-diagonal elements indicate between-muscle MCR comparison.

##### 3.2.3.2 MCR standard topography parameters (COGs and hotspots)

Considering that the MNI variant of ICC for COGs and hotspots might be more suitable when using data from many subjects (due to a smaller data variance because of the co-registration to a common brain template), we demonstrate here also the results in the MNI space (Figure 8). No muscle-specificity is observed for COGs or hotspots, either in the MNI or in the individual data.

### 3.3. Multiple-muscle MCRs interaction (within-limb TMS somatotopy) reliability

#### 3.3.1. Relative reliability

Reliability of the shifts between either COGs, or hotspots of the different muscle MCRs was poor (Table 3). At the same time, relative reliability of the areas and volumes overlaps of MCRs of different muscles was good to excellent (ICC from 0.8 to 0.9), except for the APB-ADM area and volume overlaps (ICC of which still corresponds to moderate reliability) (Table 3). ICC of EMDs between the different muscle MCR EPs was smaller (from 0.4 to 0.56), but still higher than that for COGs and hotspots shifts (Table 3).

**TABLE 3.** Relative (ICC) and absolute (SDC) reliability of the within-limb somatotopy parameters (in the individual space): COGs and hotspots’ shifts between different muscles MCRs, MCR areas and volumes overlaps, and EMDs between different muscle MCR EPs.

| Parameter, muscle | ICC | SEMeas | SDC |  |  |  |
| --- | --- | --- | --- | --- | --- | --- |
|  |  |  |  |  |  | Group<br>(n=18) |
|  |  |  | Individual |  |  |  |
|  |  |  | 68% CI | 90% CI | 95% CI | 95% CI |
| Overlap area |  |  |  |  |  |  |
| APB-ADM | 0.53 | 0.12 | 0.18 | 0.29 | 0.34 | 0.08 |
| ADM-EDC | 0.90 | 0.06 | 0.09 | 0.15 | 0.17 | 0.04 |
| APB-EDC | 0.83 | 0.07 | 0.10 | 0.16 | 0.19 | 0.05 |
| Overlap volume |  |  |  |  |  |  |
| APB-ADM | 0.48 | 0.12 | 0.16 | 0.27 | 0.32 | 0.08 |
| ADM-EDC | 0.81 | 0.08 | 0.12 | 0.19 | 0.23 | 0.05 |
| APB-EDC | 0.81 | 0.06 | 0.09 | 0.15 | 0.18 | 0.04 |
| Inter-muscle EMD, % |  |  |  |  |  |  |
| APB-ADM | 0.41 | 2.75 | 3.89 | 6.43 | 7.63 | 1.80 |
| ADM-EDC | 0.56 | 2.64 | 3.74 | 6.17 | 7.33 | 1.73 |
| APB-EDC | 0.40 | 2.71 | 3.84 | 6.33 | 7.52 | 1.77 |
| COG x, mm |  |  |  |  |  |  |
| APB-ADM | 0.15 | 1.38 | 1.95 | 3.22 | 3.83 | 0.92 |
| ADM-EDC | -0.16 | 1.60 | 2.26 | 3.73 | 4.43 | 1.08 |
| APB-EDC | -0.02 | 1.68 | 2.37 | 3.92 | 4.65 | 1.13 |

| COG y, mm |  |  |  |  |  |  |
| --- | --- | --- | --- | --- | --- | --- |
| APB-ADM | 0.35 | 1.29 | 1.83 | 3.01 | 3.58 | 0.87 |
| ADM-EDC | 0.20 | 1.16 | 1.64 | 2.71 | 3.21 | 0.78 |
| APB-EDC | 0.41 | 1.21 | 1.71 | 2.82 | 3.35 | 0.81 |
| hotspot x, mm |  |  |  |  |  |  |
| APB-ADM | 0.04 | 2.89 | 4.09 | 6.75 | 8.02 | 1.94 |
| ADM-EDC | 0.35 | 2.84 | 4.02 | 6.63 | 7.87 | 1.91 |
| APB-EDC | -0.10 | 3.76 | 5.31 | 8.77 | 10.42 | 2.53 |
| hotspot y, mm |  |  |  |  |  |  |
| APB-ADM | -0.24 | 4.80 | 6.79 | 11.21 | 13.31 | 3.23 |
| ADM-EDC | -0.28 | 5.82 | 8.24 | 13.59 | 16.15 | 3.92 |
| APB-EDC | -0.30 | 6.13 | 8.66 | 14.29 | 16.98 | 4.12 |

SEMeas and SDC values are represented in grey when ICC values are low.
CI - confidence interval.

#### 3.3.2 Absolute reliability

SDC for area and volume overlaps is in the range 0.17-0.34 (CI 95%), while SDC for the inter-muscle EMDs was around 7.5% (CI 95%) (Table 3).

#### 3.3.3. Muscle-specificity

Based on ICC before FWER, muscle-specificity can be seen for EMDs between different muscle MCR EPs (Figure 9). Based on Spearman’s correlation, areas, and volume overlaps, and inter-muscle EMDs are muscle specific and after FWER such specificity remains for the MCR areas overlaps (Figure 9).

**FIGURE 9.**
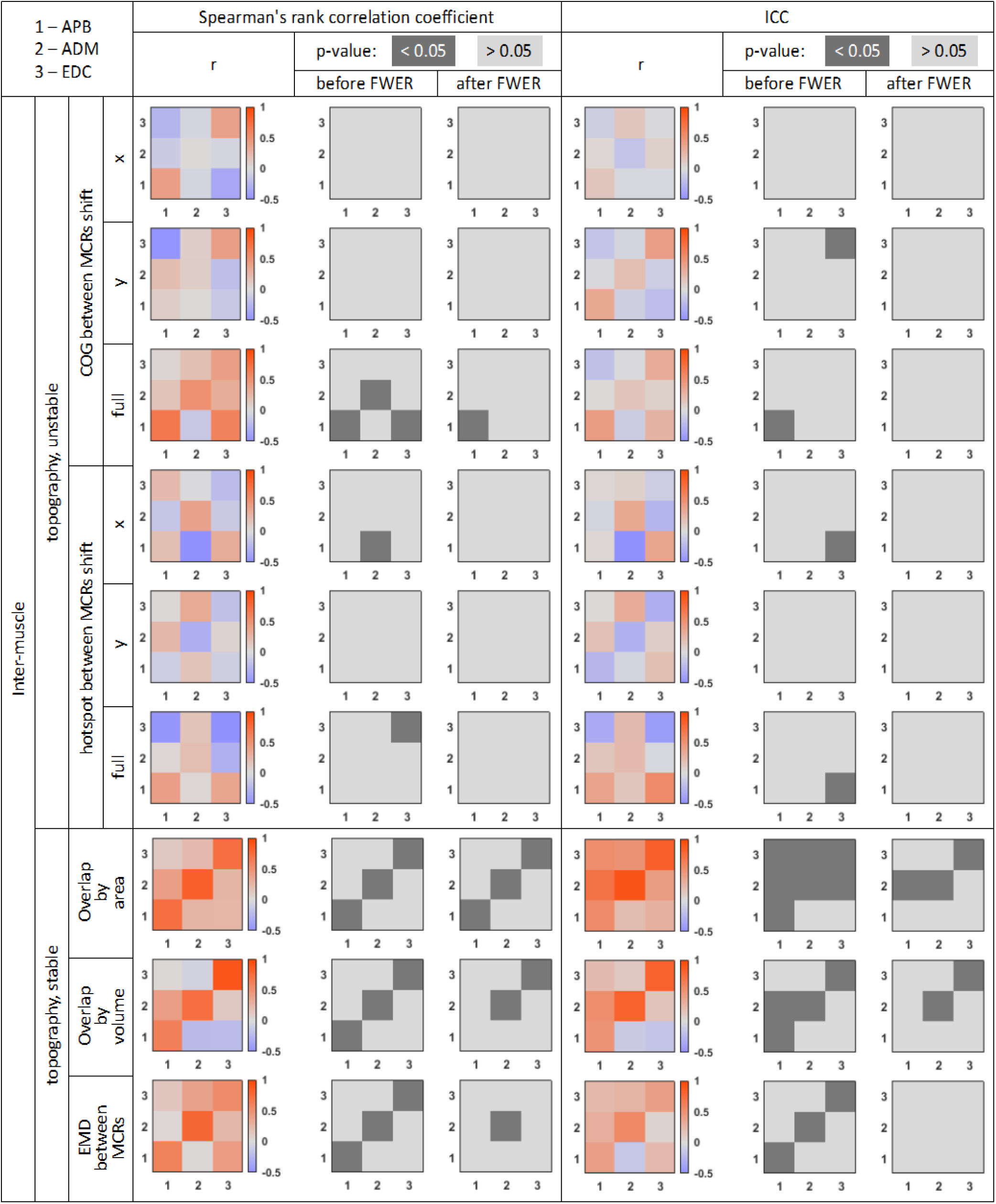
Correlation matrices for inter-muscle topography parameters in the individual space (for different muscle MCRs’ pairs: 1 – APB-ADM, 2 – ADM-EDC, 3 – APB-EDC) across days. Diagonal elements in the matrix indicate within-muscle MCR comparison, while off-diagonal elements indicate between-muscle MCR comparison.

## 4. Discussion

In this study we investigated the within-limb somatotopy gradient and the reliability of MCRs of three hand muscles in a homogenous group of healthy male volunteers, using a comprehensive grid-based sulcus-informed nTMS motor mapping. The main findings of the study are listed below:

We observed a somatotopy gradient for the hand muscle MCRs reflected by APB MCR COGs being more lateral compared to other muscles’ MCR COGs.

We demonstrated that the most commonly used metrics of MCRs, such as areas, volumes, COGs and hotspots, while having overall high relative reliability, are generally characterized by low absolute reliability.

For within-limb TMS somatotopy, the most common metrics such as the shifts between different muscles’ MCR COGs and hotspots have poor relative reliability and are not muscle-specific. While overlaps and, to a lesser extent, EMDs between different muscle MCR EPs, are more reliable and tend to demonstrate muscle-specificity.

Further we discuss each of these points in detail.

### 4.1 Somatotopy gradient

The existence of a robust within-limb somatotopy in the primary motor cortex (M1), obtained with TMS, is still debated, although “homunculus-like” TMS motor cortical maps were described from the earliest years of TMS mapping (Gentner & Classen, 2006; Metman, Bellevich, Jones, Barber, & Streletz, 1993). A commonly reported output for different muscles’ TMS MCRs interaction is the distances/shifts between the COGs (Dubbioso, Raffin, Karabanov, Thielscher, & Siebner, 2017; Schabrun, Stinear, Byblow, & Ridding, 2009; Tyč, Boyadjian, Allam, & Brasil-Neto, 2012) or hotspots (Bashir et al., 2013; Lüdemann-Podubecká & Nowak, 2016) of these different muscles’ MCRs. To the best of our knowledge, the clearest recent demonstration of the somatotopy gradient of the hand TMS MCRs has been made using “along-sulcus” TMS mapping, when one investigates MEPs produced by stimulation only along the central sulcus, taking into account its shape (Dubbioso et al., 2017; Raffin et al., 2015; Raffin & Siebner, 2019). We aimed to confirm and extend such a somatotopy gradient among the investigated hand muscle MCR COGs using a large grid approach. We observed that APB MCR COGs are more lateral compared to ADM and EDC MCR COGs. Interestingly, in two of our participants all three muscle MCRs were shifted rostrally, so it was impossible to trace MEPs along the central sulcus in a substantial part of the “hand knob” (see example in Figure 5). In this case we believe that such placement of points might correspond to a “dorsal premotor” sub-type of TMS MCRs, as opposed to the “M1” sub-type (Dubbioso, Sørensen, Thielscher, & Siebner, 2019). This supports further the notion that in order to obtain a comprehensive motor mapping in a variety of subjects one should use an extended grid to capture individual profiles of MCRs. Considering M1 division to the “new” caudal M1 (having more monosynaptic connections with the spinal motoneurons) and the “old” rostral M1 (Lemon, 2019; Rathelot & Strick, 2006) we investigated whether the “along-sulcus” MCRs (where presumably mostly “new” M1) would be more segregated compared to the “whole” MCRs. On the contrary, we found that the mediolateral somatotopy gradient is greater for the “whole” TMS mapping approach compared to the “along-sulcus” one. So, at least for our data, it appears that additional features relating to whole M1 TMS mapping do not mask the within-limb TMS somatotopy compared to the along-sulcus-only approach and might in fact provide additional topographic aspects.

We also checked the COG somatotopy gradient for the MNI version of the MCRs and showed that APB MCR is again the most lateral one. Comparing MNI MCRs across subjects, we found that the ADM MCR area tends to be smaller than the APB and EDC MCR areas, and for EDC this difference is significant. While in the “classical homunculus” the forearm representation is not only more medial but also smaller than the hand one, such a direct “homunculus” interpretation might be speculative (Nazarova & Blagovechtchenski, 2015; Schieber, 2001) and is not completely accurate even for the very same data from the original Penfield & Boldrey article, as shown in the recent re-analysis (Catani, 2017). For TMS mapping, to our knowledge, no clear difference has been reported for the sizes of the proximal upper limb muscle MCRs versus intrinsic hand muscle MCRs (Devanne et al., 2006). Considering that MNI normalization, being routine for many brain mapping techniques such as fMRI or magnetoencephalography is still not a common procedure for TMS mapping (see examples of TMS data MNI normalization (Grab et al., 2018; Kraus & Gharabaghi, 2016; Niskanen et al., 2010; Weiss et al., 2013), we believe that this confirmation of the within-hand TMS somatotopy gradient and MCR size differences in MNI space may be useful for future studies using multiple muscle TMS mapping across subjects. There is a possibility that the difference between EDC and ADM TMS MCRs is partly artificial because TMS mapping was done using 110% of APB RMT, while it was reported that the degree to which the MCR area depends on stimulation intensity may vary among muscles (Thordstein, Saar, Pegenius, & Elam, 2013). However, we believe that this cannot be a major cause of the MCR area differences because there is no clear dissimilarity among distal muscle TMS motor thresholds in healthy subjects (Ziemann, Ilić, Alle, & Meintzschel, 2004). Importantly, the difference between EDC and ADM MCRs is not prone to cross-talk between muscles, because they are located far from each other in the upper limb (Selvanayagam, Riek, & Carroll, 2012).

### 4.2 Reliability of single-muscle MCRs

A good understanding of TMS MCR reliability ranges is crucial for the TMS mapping application in longitudinal studies for the assessment of motor cortex plasticity. Here we investigated both relative reliability, allowing stratifying subjects/patients based on their MCRs, and absolute reliability, reflecting the minimal change of TMS MCR parameters, which may be traced with the current TMS mapping approach.

#### 4.2.1 Relative reliability

In the early TMS mapping studies without navigation, the relative reliability of the MCR parameters was shown to be primarily high (Malcolm et al., 2006; Plowman-Prine, Triggs, Malcolm, & Rosenbek, 2008; Wolf et al., 2004). These observations were further confirmed using non-individual brain nTMS (Cavaleri, Schabrun, & Chipchase, 2018; Jonker et al., 2019; Sankarasubramanian et al., 2015; van de Ruit, Perenboom, & Grey, 2015), as well as individual brain nTMS (Forster, Limbart, Seifert, & Senft, 2014; Kraus & Gharabaghi, 2016; Sollmann et al., 2013; Weiss et al., 2013; Zdunczyk, Fleischmann, Schulz, Vajkoczy, & Picht, 2013). The problem is that only COG location reliability was always reported in TMS mapping papers, while the inclusion of other parameters was inconsistent. Using individual brain nTMS mapping it was shown that MCR parameters such as areas and volumes may be less reliable compared to COGs and mean MEP per MCR (Kraus & Gharabaghi, 2016). Previously, the calculations of MCR parameters were based on variable custom-made scripts, so we aimed at accessing the reliability of MCR parameters obtained using TMSmap software (Novikov et al., 2018), which we suggested as a possible unifying approach for the fast and easy quantitative assessment of TMS motor mapping results. We observed high ICC values for all the investigated standard metrics of TMS MCRs (areas, volumes, mean MEPs, hotspot and COG locations), with the highest ICC values for COGs and hotspots (ICC>=0.9). However, for the normalized MNI data, the ICC for hotspots and COGs was substantially lower because of the lower data variance among the brains after normalization, but still generally remained good.

#### 4.2.2. Absolute reliability

Absolute reliability (which may be reported by SEM, SDC or limits of agreement) is crucial for the method’s use on an individual level in clinical practice or sport (Atkinson & Nevill, 1998; Beaulieu, Flamand, et al., 2017). The absolute reliability of TMS motor mapping results has been described in the literature less frequently than the relative one (Jonker et al., 2019; Ngomo, Leonard, Moffet, & Mercier, 2012; Potter-Baker et al., 2016; van de Ruit et al., 2015). To the best of our knowledge, previous TMS motor mapping studies reporting classic absolute reliability metrics, were performed using non-individual MRI navigation, except one study (Ngomo et al., 2012). Yet the importance of considering individual anatomy in TMS motor mapping has been emphasized (Bashir et al., 2013; Raffin et al., 2015). When absolute reliability was reported, it was recommended to be interpreted “with caution on an individual level” (Jonker et al., 2019). In our study we provide a comprehensive range of SDC values for different MCR parameters for three hand muscles. We can also interpret SDC as being generally sizeable: a particularly thought provoking SDC was found for the hotspot coordinates (around 1 cm), it also corresponds with a recent finding that hotspot may be considered as an area rather than a point taking into account MEPs variability and EF-spread (Reijonen et al., 2020). Hotspot is a crucial parameter for TMS motor mapping studies (Bashir et al., 2013; Rossini et al., 2015; Sollmann et al., 2013), where shifts of just several millimeters between hotspots of different MCRs are reported (Bashir et al., 2013). Hotspot is also highly relevant for a wide range of TMS applications with repetitive sessions of stimulation, including therapeutic TMS. Thus, we suggest that for longitudinal TMS design (either for diagnostic or therapeutic purposes) it is important to recheck the hotspot location with a roughly 1 cm radius, at least every several days.

#### 4.2.3. Muscle-specificity

We were interested in the possibility of reliably tracing muscle-specific features of different muscle MCRs. Thus, we introduced a term “muscle-specificity” of the TMS MCR parameter, which we defined as the possibility to predict a given muscle among others (but only when using the data across subjects), using a certain MCR parameter. We observed that among standard parameters of single-muscle MCRs only mean MEP tended to be muscle-specific. Additionally, we investigated the reliability of a more novel parameter of the EP, which we previously implemented in TMSmap software (Novikov et al., 2018), similar to the EP probed with the along-sulcus mapping approach (Raffin et al., 2015; Raffin & Siebner, 2019). We showed that the difference between the EPs, reflected in the EMD values (Novikov et al., 2018; Rubner et al., 1998) of the same muscle between days being smaller than for the different muscles MCR EPs between days. We believe that the idea of the MCR EP is important because it relates to the key phenomenon of motor cortex organization – convergence (Schieber, 2001), manifested in the well-known fact that different neurons in M1 may be involved in a given muscle activity depending on the goal/motor task or other conditions (Capaday, Ethier, Van Vreeswijk, & Darling, 2013). Moreover, using fMRI it has been shown that a single-muscle MCR is not homogeneous in its brain connectivity patterns (Smith et al., 2017). A similar approach of discrete peak calculation in an MCR was proposed recently (Elgueta-Cancino, Marinovic, Jull, & Hodges, 2019; Massé-Alarie, Bergin, Schneider, Schabrun, & Hodges, 2017) but it has been already shown to be less reliable than the standard TMS MCR metrics (Cavaleri et al., 2018).We also consider EPs comparison using EMD to be more direct than discrete peak calculation because while using EMD one does not need to define an arbitrary threshold level of the peak, and it is possible to account for the complex two-dimensional shape of the MCR. However, further validation of the MCR EP comparison using EMD in an interventional longitudinal TMS mapping studies is necessary.

#### 4.2.4. Reliability of the MCR overlaps

While we confirmed the somatotopy gradient between APB MCR versus ADM and EDC MCRs, using COG comparison, the relative reliability of the shifts between different muscle MCR COGs and hotspots is very poor and these metrics are not muscle-specific. We evaluated MCR overlap reliability as another parameter of the within-limb somatotopy. Extensive overlaps among different muscle MCRs were described from the early TMS studies (Devanne et al., 2006; Melgari, Pasqualetti, Pauri, & Rossini, 2008; Wassermann et al., 1993). Such overlaps reflect yet another key principle of M1 organization – its divergence, meaning that the output of a single M1 neuron may reach multiple spinal motoneurons, resulting in the activation of different limb segments (Schieber, 2001). It was demonstrated both non-invasively by TMS (Gerachshenko, Rymer, & Stinear, 2008) and invasively by microstimulation (Graziano, 2006), that the stimulation of the same cortical point may evoke activity in different muscles depending on a number of factors such as limb position (Graziano, 2006), anticipated movement (Gerachshenko et al., 2008; Uehara, Morishita, Kubota, & Funase, 2013) etc. Moreover, it was recently reported that in pathological cases, such as a tetraplegia, the “hand knob” in the precentral gyrus may be tuned not only to other upper limb segments but to the entire body (Willett et al., 2020). There is also some proof that the extent of TMS MCR overlaps may change in pathological conditions like dystonia or chronic pain (Schabrun, Hodges, Vicenzino, Jones, & Chipchase, 2015; Schabrun et al., 2009). We observed that the overlaps between different muscle MCRs, and, to a lesser extent, EMDs between the different muscle MCRs, were more reliable and tended to be muscle-specific. To our knowledge, this is the first demonstration of the reliability of the inter-muscle interactions for multiple muscle TMS motor mapping using individual MRI data. Another hypotheses about MCR overlaps is that they may be a neural substrate underlying muscle synergies (Beaulieu, Flamand, et al., 2017; Capaday et al., 2013; Pitkänen et al., 2017; Schabrun et al., 2015, 2009). While, for instance, it has been discussed that the primary source of after-stroke pathological synergies may be in the medulla (Karbasforoushan, Cohen-Adad, & Dewald, 2019; McPherson et al., 2018; Zaaimi, Dean, & Baker, 2018), the MCR overlap might at least partly represent a cortical manifestation for this phenomenon (Giszter, 2015; Huffmaster, Van Acker, Luchies, & Cheney, 2018; Klochkov, Khizhnikova, Nazarova, & Chernikova, 2017). Therefore, finding the high reliability of MCR overlaps and their muscle-specificity may be encouraging for new studies where such overlaps may be used as a traceable parameter reflecting training/rehabilitation dedicated to synergy manipulation.

## 5. Methodological considerations and future studies

An important strength of this study is the simultaneous investigation of multiple muscle MCRs. To our knowledge, this is the first work where the absolute reliability of multiple muscle MCRs was probed, using individual brain navigated TMS mapping for the whole motor cortex, in addition to the along-sulcus TMS mapping (Raffaele Dubbioso et al., 2017; Raffin et al., 2015). Another feature of this study is its robust mapping design: we used up to 350 points per subject, because the literature about the necessary number of points for TMS motor mapping is contradictory: while some parameters of MCR were reported to be reliably traced already with 60 stimuli (van de Ruit et al., 2015), there have also been indications that accuracy of MCR parameters continued to increase without saturation up to a much higher number of stimuli (Nazarova et al., 2018; Sinitsyn et al., 2019). Another methodological strength of our study is the homogenous group of male participants with no special motor skills. Yet it may also be viewed as a limitation, meaning that in a more diverse multi-gender population the reliability of motor parameters may be less pronounced, thus a separate reliability study in females, taking, for instance, into account the menstrual cycle, may be needed. Another issue is that we probed only hand muscles and thus the reliability for distal versus proximal upper limb muscle MCRs is left for further investigation. Also, we did not employ any motor task to assess motor abilities, thus, the question of the functional relevance of TMS within-limb somatotopy and its reliability should be investigated in the future. Yet other technical limitation is that when tracing several muscle MCRs, the characteristics of the MCR parameters distribution (such as normality or homoscedasticity) may vary across muscles, so, for instance, the question of the logarithmic transformation for the parameters has to be carefully considered.

One general limitation applicable not only to our work but to all TMS studies, is the spatial resolution of TMS. We assume that the simultaneous activation of several muscles by TMS at one point is a result of several key factors: (1) the non-focality of TMS itself, where a spread of induced electric field leads to the activation of a considerable cortical area (Wassermann et al., 2008); (2) the primarily indirect TMS effect on the pyramidal neurons (Seo, Schaworonkow, Jun, & Triesch, 2016; Spampinato, 2020); (3) the co-location of the cortical neuronal populations, innervating different spinal motoneurons pools (Capaday et al., 2013; Schieber, 2001); and (4) cross-talk between the muscles at the peripheral level when using surface EMG (Selvanayagam et al., 2012). TMS focality depends on the coil configuration and on the intensity, shape and direction of the pulse (Koponen, Nieminen, Mutanen, Stenroos, & Ilmoniemi, 2017; Rossini et al., 2015; Sommer et al., 2006, 2018; Tugin et al., 2020). We utilized the figure-of-eight coil used for nTMS presurgical mapping (Krieg et al., 2017) and a biphasic pulse shape, allowing to use less intensity (Raffin et al., 2015). We used a biphasic pulse shape because it is the most common in TMS mapping studies with patients (Lüdemann-Podubecká & Nowak, 2016; Takahashi, Vajkoczy, & Picht, 2013). However, considering that a biphasic pulse may activate two distinct neuronal pools (Sommer et al., 2018); it may be informative to investigate how the reliability of the within-limb TMS somatotopy differs when using monophasic current configurations.

## 6. Conclusions

Until now the use of nTMS mapping for longitudinal purposes in fundamental and clinical studies remains challenging, especially when multiple MCRs are probed. In this work we confirmed the existence of the somatotopy gradient for the hand MCR COGs for the whole motor cortex nTMS mapping in addition for the along-sulcus TMS mapping approach. Adding to previous reliability studies, we confirmed a high relative reliability of the standard MCR parameters for three hand muscles and present a range of SDC values for the MCR parameters. We provide novel evidence that inter-muscle MCR interaction can be reliably traced using MCR area overlaps, while shifts between the COGs and hotspots of different muscle MCRs are not suitable for this purpose. Our work has also a practical perspective – a high reliability of MCR overlaps allows us to suggest them as a possible cortical biomarker for tracking the neuronal changes associated with the training/rehabilitation aimed at the modifications of muscle synergies.

## Acknowledgments

nTMS data were obtained in the Research Center of Neurology, Moscow while MN was working there in the years 2012-2016. Authors are deeply grateful to prof. M.A. Piradov for his administrative support during nTMS data collection. Authors are also grateful to I. Gusarovas and D. Pozdeeva for software testing and help with the experiments. We are also grateful to Dr. Eini Niskanen for her recommendations considering MNI data normalization. The work of MN was partly supported by Russian Science Foundation grant 19-75-00104.

